# PA-X is an avian virulence factor in H9N2 avian influenza virus

**DOI:** 10.1101/2020.05.25.114876

**Authors:** Anabel L. Clements, Thomas P. Peacock, Joshua E. Sealy, Saira Hussain, Jean-Remy Sadeyen, Holly Shelton, Paul Digard, Munir Iqbal

**Author notes:** The Francis Crick Institute, London, UK, NW1 1AT. Corresponding author: Munir Iqbal.

## Abstract

Influenza A viruses encode several accessory proteins that have host- and strain-specific effects on virulence and replication. The accessory protein PA-X is expressed due to a ribosomal frameshift during translation of the PA gene. Depending on the particular combination of virus strain and host species, PA-X has been described as either acting to either reduce or increase virulence and/or virus replication. In this study, we set out to investigate the role PA-X plays in H9N2 avian influenza viruses, focussing particularly on the natural avian host, chickens. We found H9N2 PA-X induced robust host shutoff in both mammalian and avian cells and increased replication in mammalian, but not avian cells. We further showed that PA-X affected embryonic lethality *in ovo* and led to more rapid viral shedding and widespread organ dissemination *in vivo* in chickens. Overall, we conclude PA-X may act as a virulence factor for H9N2 viruses in chickens, allowing faster replication and wider organ tropism.

## Introduction

Influenza A viruses (IAV) have segmented negative-sense RNA genomes encoding 10 core proteins and a variable number of strain-specific accessory proteins (Vasin et al., 2014). Due to its small genome size and nuclear replication, IAV has evolved a number of ways to increase its protein coding capacity including the use of splice variants (M2, NEP and the more recently discovered M42, PB2-S1 and NS3 proteins), encoding multiple open reading frames (ORFs), both nested and overlapping on a single gene segment (e.g. PB1-F2, PB1-N40, NA43) and ribosomal frameshifts leading to multi-ORF fusion proteins (PA-X) (Wise et al., 2012, Wise et al., 2009, Yamayoshi et al., 2016, Selman et al., 2012, Chen et al., 2001, Machkovech et al., 2019)

Influenza A virus segment 3 primarily encodes the polymerase acidic protein (PA). PA is an integral part of the influenza virus RNA-dependent RNA polymerase (RdRp) and contains two functional domains, an N-terminal endonuclease (endo) domain, responsible for cleaving host capped RNAs used to prime viral transcription, and a C-terminal domain, associated with the core of the RdRp (Te Velthuis and Fodor, 2016). The PA endo domain contains a conserved ribosomal frameshift site – a rare arginine codon facilitates ribosomal stalling, which is followed (at low level) by ribosomal slippage into a +1 open reading frame (ORF), facilitated by an upstream UUU codon that allows realignment (Firth et al., 2012, Jagger et al., 2012). The resulting protein, PA-X, is a fusion between the first 191 amino acids of PA (the endo domain) and up to 61 amino acids translated from the X-ORF in the +1 reading frame. PA-X has been shown to mediate degradation of host cell mRNAs and disruption of host mRNA processing, leading to host cell shutoff and a dampened innate immune response (Jagger et al., 2012, Gaucherand et al., 2019, Khaperskyy et al., 2016, Oishi et al., 2015, Lee et al., 2017, Hayashi et al., 2016, Desmet et al., 2013).

H9N2 avian influenza viruses are enzootic through much of Asia, the Middle East and Africa (Peacock et al., 2019). H9N2 viruses cause massive economic damage due to their impact on poultry production systems, causing moderate morbidity and mortality, especially in the context of viral or bacterial coinfection. Additionally, H9N2 viruses pose a direct zoonotic threat to humans and are considered viruses with pandemic potential (Peacock et al., 2019, Song and Qin, 2020). As well as being zoonotic threats in their own right, H9N2 viruses have contributed polymerase genes to multiple zoonotic avian influenza viruses, including epidemic H7N9 (Liu et al., 2013).

Several studies have investigated the role of PA-X on virulence and replication of avian influenza viruses in mammalian and avian hosts. There is little consensus between these studies and there appears to be virus strain- and host species-specific differences in whether PA-X expression increased or decreased replication, virulence and transmissibility of IAV (Gao et al., 2015a, Gao et al., 2015b, Gao et al., 2015c, Jagger et al., 2012, Gong et al., 2017, Hussain et al., 2019, Lee et al., 2017, Sun et al., 2020, Hu et al., 2015, Rigby et al., 2019, Ma et al., 2019). In H9N2 viruses, expression of full length PA-X has been shown to be a virulence factor in mammalian infection systems (Gao et al., 2015c, Gao et al., 2015a). However, it is unclear what role PA-X has in these viruses in their natural chicken hosts.

In this study we set out to investigate the role of PA-X in a contemporary G1-lineage H9N2 virus, typical of viruses circulating in the Middle East and South Asia (Peacock et al., 2019). We found that the H9N2 virus expressed a PA-X capable of robust host shutoff which correlated with PA-X expression. Removing PA-X expression decreased viral replication in mammalian, but not avian cell culture systems, although it did reduce embryonic lethality *in ovo*. *In vivo*, in the natural chicken host, ablation of of PA-X expression led to delayed shedding and restricted viral dissemination. Overall, our results suggest PA-X may be an H9N2 virulence factor in the natural avian host, allowing faster replication and increased visceral organ tropism.

## Results

### Generation of H9N2 viruses with altered PA-X expression

To investigate the role of PA-X in an H9N2 background we generated a panel of mutants in the background of a contemporary G1-lineage H9N2 virus, A/chicken/Pakistan/UDL-01/2008 (UDL-01), typical of viruses still circulating in South Asia and the Middle East (Iqbal et al., 2009, Sealy et al., 2018). Mutants were generated (in cDNA copies of segment 3 cloned into a reverse genetics plasmid) using a previously validated approach, with the frameshift site mutated in such a way that ribosomal slippage should be inhibited (FS); additionally, a panel of truncated PA-X mutants with premature termination codons (PTC) spaced throughout the X-ORF, were made (Table 1, Figure 1A) (Jagger et al., 2012, Hussain et al., 2019). All mutations were synonymous in the coding sequence of PA.

**Table 1.** Summary of PA-X mutants made in this study

| Mutation name | Original nt | Mutated nt | nt position(s) | PA-X Amino acid change |
| --- | --- | --- | --- | --- |
| **FS** | TCC TTT CGT | AGC TTC AGA | 568,569,573,574,576 |  |
| **PTC1** | C | A | 621 | I207STOP |
| **PTC2** | C, C, G | A, G, A | 634,636,642 | R212STOP, L214STOP |
| **PTC3** | T | A | 678 | L226STOP |
| **PTC4** | G | A | 699 | V233STOP |

**Figure 1.**
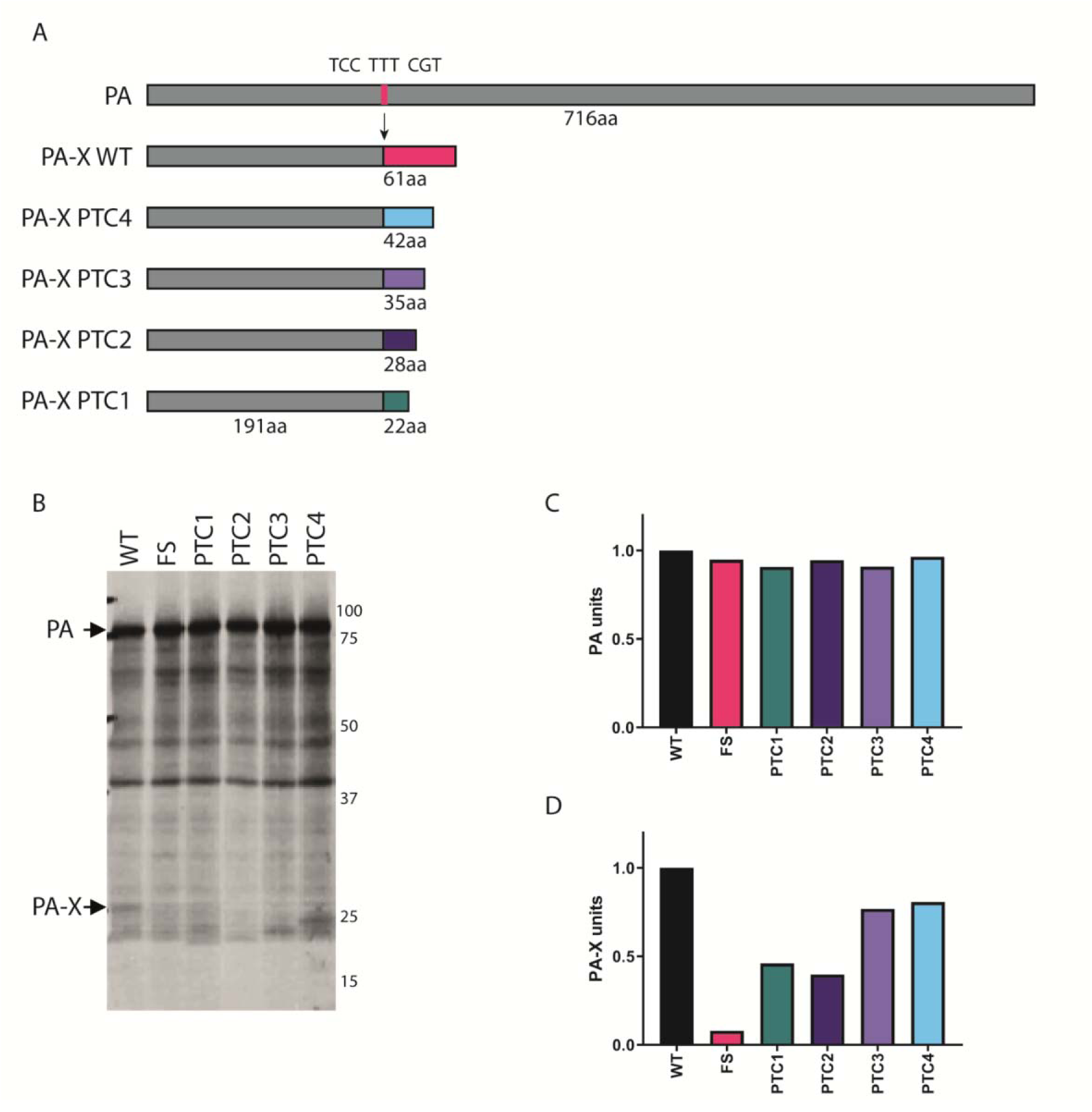
Generation and validation of H9N2 viruses with mutant PA-X proteins. A panel of mutations were made within UDL-01 segment 3 that altered PA-X expression. (A) Location of mutations within frameshift site and X-ORF. Dark grey rectangle represents PA, light grey rectangle represents X-ORF. Pink line represents location of the frameshift site (FS), coloured lines represents location of PTC mutations. (B) Coupled in vitro transcription-translation reactions radiolabelled with ^35^S-methionine were carried out using the TnT rabbit reticulocyte lysate system and protein products analysed using SDS-PAGE and autoradiography. PA and PA-X expression are marked via black arrows. (C) quantification of the AUC of the densitometry analysis of the PA band using ImageJ analysis software. (D) quantification of the AUC of the densitometry analysis of the PA-X band using ImageJ analysis software.

To confirm ablation of PA-X expression and/or validate the truncations, *in vitro* transcription/translation assays in rabbit reticulocyte lysate were carried out directly from the plasmids. Comparable levels of PA expression were seen across every construct, confirming the PA-X mutations did not affect PA expression (Figure 1B, C). As expected (Jagger et al., 2012, Hussain et al., 2019, Lee et al., 2017), PA-X expression was drastically reduced in the FS mutant (Figure 1B, D). Upon X-ORF truncation, a ladder effect where the size of PA-X was progressively decreased by the PTC mutations could be seen, while both PTC1 and PTC2 also showed a reduction of PA-X expression. Overall, these results indicate that a previously used strategy for altering PA-X expression is also successful in this H9N2 virus background.

### A PA-X frameshift mutation abrogates host cell shutoff activity

PA-X plays a key role in IAV host cell shutoff, therefore the ability of the H9N2 PA-X variants to repress cellular gene expression was tested using transfected segment 3 plasmids in β-galactosidase (β-gal) reporter assays. HEK 293Ts or DF-1 cells were transfected with the β-gal plasmid along with segment 3 plasmids or an empty vector control, β-gal accumulation was measured by enzyme assays 48 hours later and normalised to empty vector. UDL-01 WT PA-X but not the A/Puerto Rico/8/34 (PR8) PA-X showed robust repression of β-gal expression in both cell types, and introduction of the FS mutant ablated the UDL-01 activity (Figure 2A, B), indicating that the shutoff activity of segment 3 is dependent on PA-X expression but varies according to IAV strain, as previously shown (Hussain et al., 2019, Jagger et al., 2012, Desmet et al., 2013). When shut-off activity of the UDL-01 PTC mutants was tested in 293T cells, PTC1 and PTC2 showed a minor and non-statistically significant reduction in host shutoff activity compared to UDL-01 WT while PTC3 and PTC4 had no apparent effect. Thus UDL-01 encodes an active PA-X polypeptide, whose shutoff function does not strongly depend on the full X-ORF sequence.

**Figure 2.**
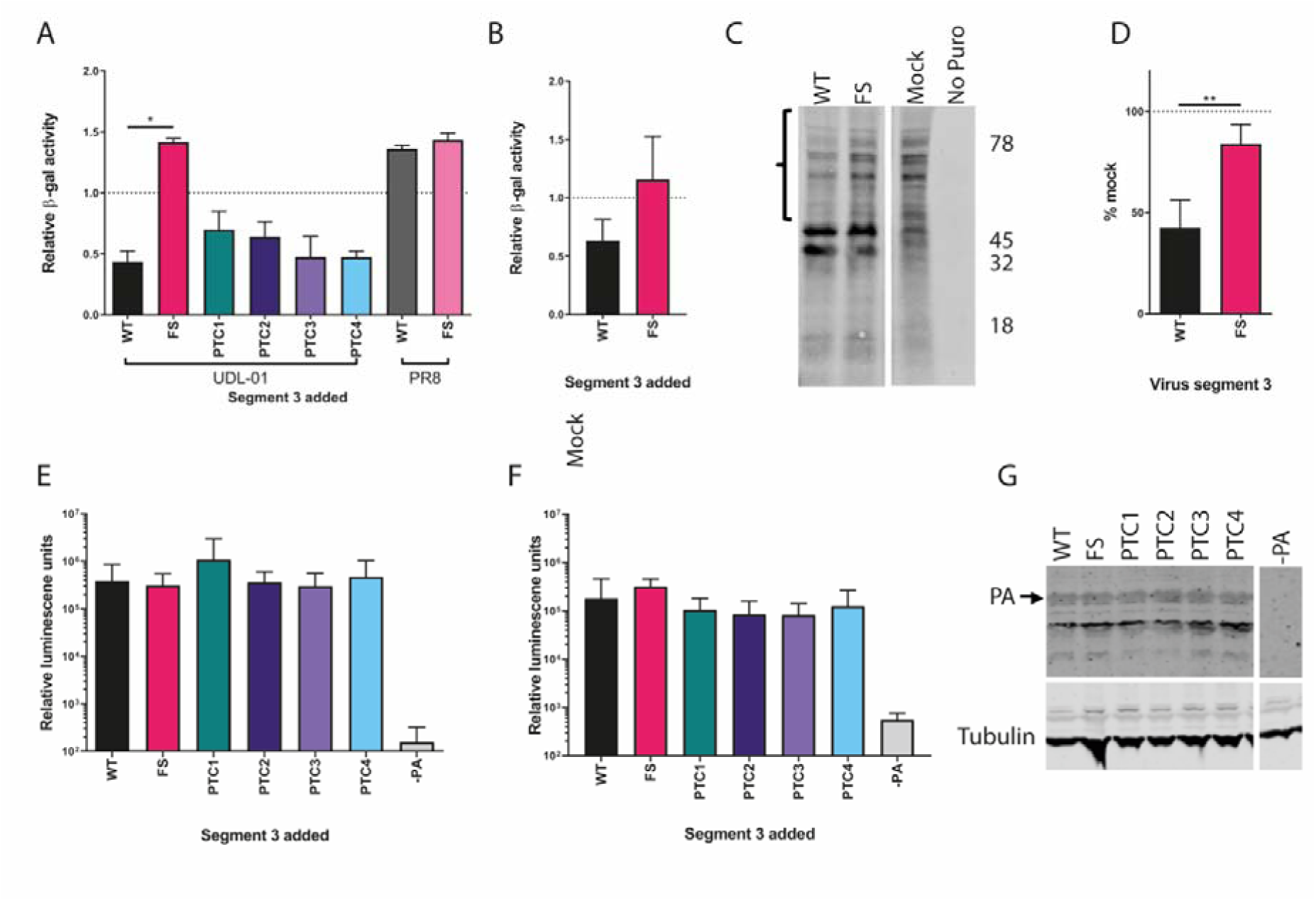
Ablation of PA-X expression leads to a loss of host cell shutoff but has no effect on virus polymerase activity. (A) 293T cells or (B) DF-1 cells were transfected with a β-gal reporter plasmid alongside the indicated segment 3 expression plasmids with or without PA-X mutations. 48 hours post-transfection cells were lysed and levels of β-gal enzymatic activity assessed by colorimetric assay. Results were normalised to an empty vector control. Graphs represent the average +/− SD of 3 independent experiments. Statistical significance was determined by one-way ANOVA with multiple comparisons (A) or unpaired T-test (B). (C, D) MDCK cells were infected with a high MOI (5) using viruses expressing PA-X (WT) or with PA-X expression removed (FS). Seven hours post-infection cells were pulsed with puromycin for 30 minutes and then lysed and samples run on SDS-PAGE gels. Membranes were probed for presence of puromycin. (C) Representative SDS-PAGE gel with the area quantified highlighted in brackets. The approximate position of molecular mass markers (kDa) is also indicated. (D) Quantification of the densitometry of the highlighted area (above 47 kDa) performed using ImageJ analysis software. Graph represents the average of 3 independent experiments +/− SD. Each data group is normalised to mock levels of protein synthesis. Statistics determined by unpaired T-test. (E) 293T cells or (F) DF-1 cells were transfected with the components of the polymerase complex (PB1, PB2, PA and NP) plus a vRNA mimic encoding luciferase. 48 hours post-transfection cells were lysed and luciferase levels measured. Data are the average of 3 independent experiments +/− SD. Statistics were determined by Kruskal Wallis test with multiple comparisons. (G) PA and tubulin expression levels were determined via western blot analysis of 293T cell lysates from part (E). P values for statistics throughout: *, 0.05 ≥ P > 0.01; **, 0.01 ≥ P > 0.001

To investigate whether the effect of the FS mutation on host shutoff activity of UDL-01 segment 3 seen with plasmid-based assays could be recapitulated with infectious virus, mutant viruses were generated by reverse genetics. MDCK cells were then infected with WT and FS UDL-01 viruses at a high MOI (5), and pulsed with puromycin for 30 minutes to label nascent polypeptides (Schmidt et al., 2009), before being lysed. Lysates were then run on SDS-PAGE western blotted for puromycin (Figure 2C). The region of the blot corresponding to ~ 80-50 kDa, above where a pair of virally induced protein bands were seen, was assessed by densitometry to measure host protein synthesis levels within the cell. UDL-01 WT virus reduced cellular protein synthesis by over 50% compared to uninfected cells while the FS mutant only caused <20% host shutoff (Figure 2D). These data corroborate the plasmid-based methods previously used and showed that in the context of infectious virus, UDL-01 expresses a classically active PA-X protein.

### PA-X expression does not affect H9N2 polymerase activity

Influenza PA-X has been suggested to modulate polymerase activity (Hu et al., 2015, Lee et al., 2017, Gao et al., 2015b, Gong et al., 2017), although these studies did not all agree on whether PA-X promotes or suppresses polymerase activity; the effect may be strain-as well as cell-type dependent. Therefore, we investigated the effect of PA-X expression on H9N2 polymerase activity in avian and mammalian cells. Cells were co-transfected with plasmids encoding the polymerase components and NP, alongside a viral RNA-like reporter encoding luciferase. In 293Ts, PB2 and PB1 from the mammalian adapted strain PR8 were used to overcome the restriction of avian IAV polymerase in these cells, whereas in avian DF-1 cells, the full polymerase from UDL-01 was used. Exchanging WT UDL-01 segment 3 with the different PA-X mutants had no significant effect on polymerase activity in either mammalian or avian cells (Figure 2E, F). Furthermore, upon western blotting cell lysates, no differences in PA or tubulin expression were seen, indicating that PA-X expression did not alter PA accumulation (Figure 2G).

### Viruses with abrogated PA-X expression have a minor replicative defect in mammalian but not avian cells

Although no difference in polymerase activity was seen between the different PA-X mutants, multicycle growth curves were performed in mammalian and avian systems to determine whether differences in PA-X may affect viral replication in a more biologically relevant context. PA-X removal has been shown to impact viral replication in several previous studies (Lee et al., 2017, Hu et al., 2015, Gao et al., 2015b, Gao et al., 2015c), though similarly to polymerase activity these studies tend to disagree about whether PA-X expression enhances or suppresses viral replication and here too, the effect may be virus strain and host-specific.

Initially, plaque size in MDCKs was assessed to determine if any gross replication defects could be seen in the mutants as plaque phenotype is a proxy for replicative fitness in influenza viruses. A modest, but significant reduction in plaque diameter was observed within UDL-01 when PA-X was removed or truncated up to PTC3 (Figure 3A, B). Average plaque diameters decreased from 1.7 mm to 1.2 mm, 1.25 mm, and 1.36 mm respectively.

**Figure 3.**
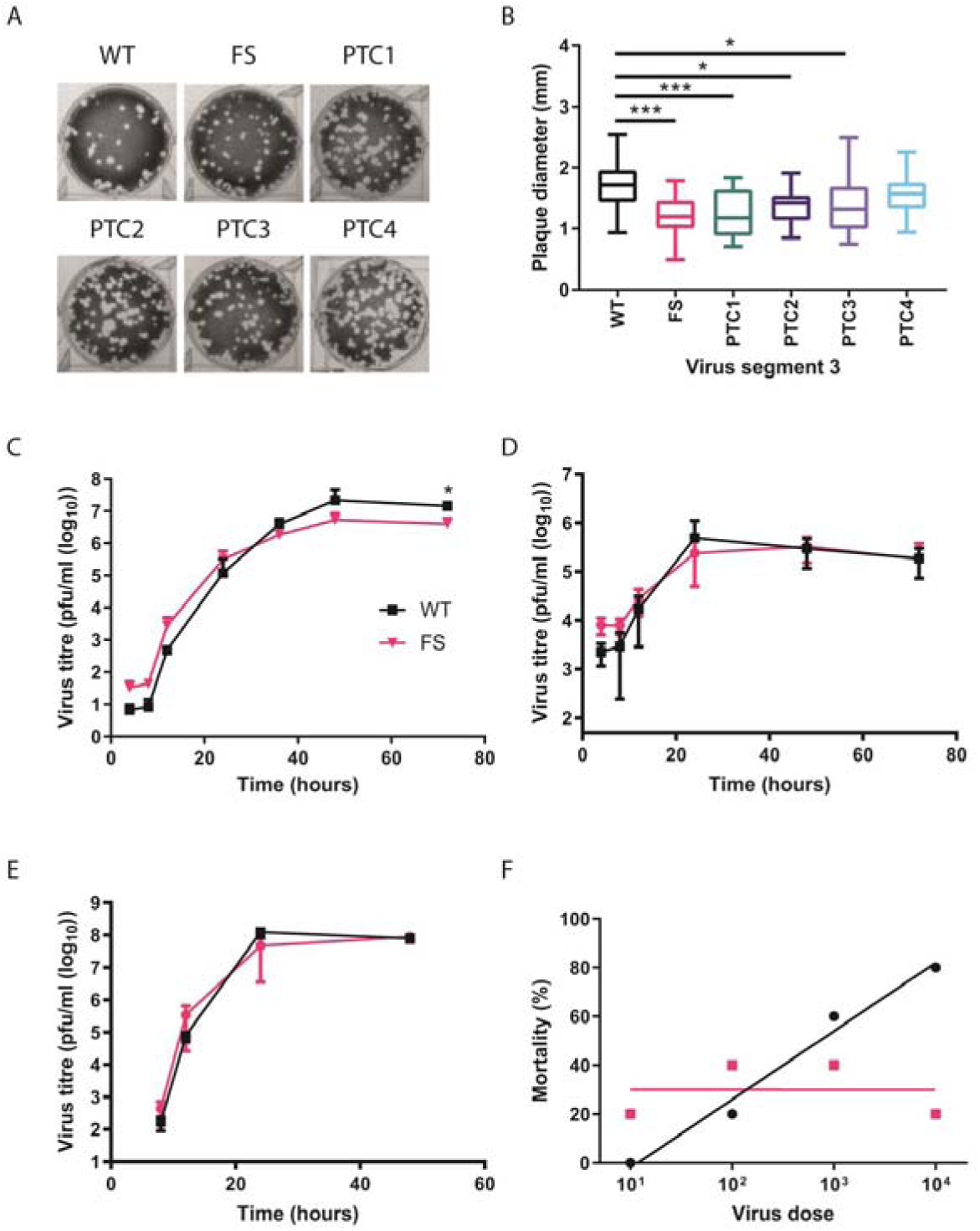
PA-X deficient viruses have a minor growth defect and lower *in ovo* mortality. (A,B) Viruses were rescued using reverse genetics and titrated under a 0.6% agarose overlay in order to ascertain the plaque phenotype. (A) After 72 hours cells were fixed and stained with 0.1% crystal violet solution and plates imaged. (B) ImageJ analysis software was used to measure the diameter of 20 plaques per virus. Graph represents average diameter of 20 plaques +/− SD. Statistical significance was determined by one-way ANOVA with multiple comparisons. (C) MDCK cells or (D) CK cells were infected with a low MOI (0.01) of WT or FS H9N2 viruses. Cell supernatants were harvested at 4, 8, 12, 24, 36, 48 and 72 hours post-infection and titrated via plaque assay. Graphs represent an average of 3 independent experiments +/− SD. (E) 10-day old fertilised hens’ eggs were infected with 100 pfu of each virus. Allantoic fluid was collected at 4, 8, 12, 24 and 48 hours post-infection and titrated via plaque assay. Graph represents an average of 5 eggs per virus per time point +/− SD. Statistics through determined by Mann-Whitney U test. (F) Embryonated hens’ eggs were infected with different doses of virus, at 84 hours post-infection, total embryo mortality after infection with the indicated viruses at each viral dose was calculated. Line represents non-linear fit of data with each data point represent % mortality at the viral dilution. P values for statistics throughout: *, 0.05 ≥ P > 0.01; ***, 0.001 ≥ P > 0.0001.

As UDL-01 FS had the largest impact on plaque diameter and host shutoff, this virus was taken forward to further viral replication kinetics experiments. MDCKs were infected at a low MOI and virus titres were assessed over a time course. UDL-01 WT and FS showed very similar growth kinetics, although UDL-01 WT displayed significantly higher titres at 72 hours post-infection compared to UDL-01 FS (Figure 3C). From 48 hours post-infection there was a trend for decreased viral titres with UDL-01 FS compared to UDL-01 WT, implying in this particular H9N2 strain PA-X expression may slightly enhance viral replication in MDCK cells.

We next performed a similar growth kinetics experiment in avian primary chicken kidney (CK) cells. In contrast to the results seen within MDCK cells there was no significant difference in viral replication between UDL-01 WT and FS at any time point, suggesting a host-specific effect (Figure 3D). To assess this further, replication kinetics were assessed in 10-day-old fertilised hens’ eggs. Eggs were inoculated with 100 pfu of each virus and allantoic fluid was harvested periodically. *In ovo*, UDL-01 WT and FS viruses did not exhibit any significant differences in viral replication throughout the course of infection (Figure 3E). Overall, the impact of mutating PA-X on the replication of H9N2 AIVs was variable and replication differences appeared to be host-dependent with PA-X playing a role on replication in mammalian, but not avian systems.

### Viruses with abrogated PA-X expression have lower embryonic lethality

PA-X expression has previously been shown to alter viral pathogenicity in animal models e.g. (Jagger et al., 2012, Gao et al., 2015b, Gao et al., 2015c, Gong et al., 2017, Hu et al., 2015, Rigby et al., 2019) as well as *in ovo* (Hussain et al., 2019). Prior to performing an *in vivo* experiment we infected 10-day-old embryonated hens’ eggs with serial dilutions of UDL-01 WT or FS mutants and assessed embryonic lethality over 84 hours, as previously described (Hussain et al., 2019). When percentage survival was plotted against viral dilution a clear difference could be seen between UDL-01 WT and FS (Figure 3F). The inoculum size of UDL-01 WT negatively affected embryo survival in a dose dependant manner as seen by the ascending trend line, whereas UDL-01 FS survival was not dose-dependent. Overall, these data were suggestive of a difference in a pathogenicity between the WT and PA-X deficient viruses.

### Viruses lacking PA-X expression have delayed shedding and reduced visceral tropism *in vivo*

As the effect of PA-X expression has yet to be assessed in H9N2 viruses in their natural host, we performed an *in vivo* experiment to assess the effect of PA-X on virus replication, transmission, tropism, pathogenicity and cytokine expression in chickens. Groups of 10 three-week-old White Leghorn (VALO breed) chickens were inoculated intranasally with 10^4^ pfu of either UDL-01 WT or FS virus (or sterile PBS). One day post-inoculation, 8 naïve contact birds were introduced into each directly infected group to assess viral transmission. Birds were swabbed daily in both buccal and cloacal cavities to determine viral shedding. Throughout the study period birds were monitored for clinical signs, however only very minimal signs were seen in any birds (data not shown). Furthermore, in both directly infected and contact birds, no culturable virus was detected from cloacal swabs. Directly infected birds in both groups showed robust buccal shedding from days 1-6, peaking at titres of over 10^4^ pfu/ml (Figure 4A). However, birds infected with UDL-01 FS showed a delayed buccal shedding profile compared to UDL-01 WT infected birds, shedding significantly less on days 1-2. By day 3 post-infection, shedding levels were comparable between groups and by day 6, an increased number of animals infected with UDL-01 FS shed compared to WT (2/7 for UDL-01 WT vs 4/6 for UDL FS). Within both groups, viral shedding was cleared by day 7 post-infection. To estimate the total amount of virus shed by each group, the area under the shedding curves (AUC) were calculated, somewhat comparable values were obtained for both viruses; 81,674 for WT UDL-01 versus 241,229 for the FS mutant. This suggested that UDL-01 FS infected birds shed more virus buccally over the course of the experiment than the UDL-01 WT infected birds.

**Figure 4.**
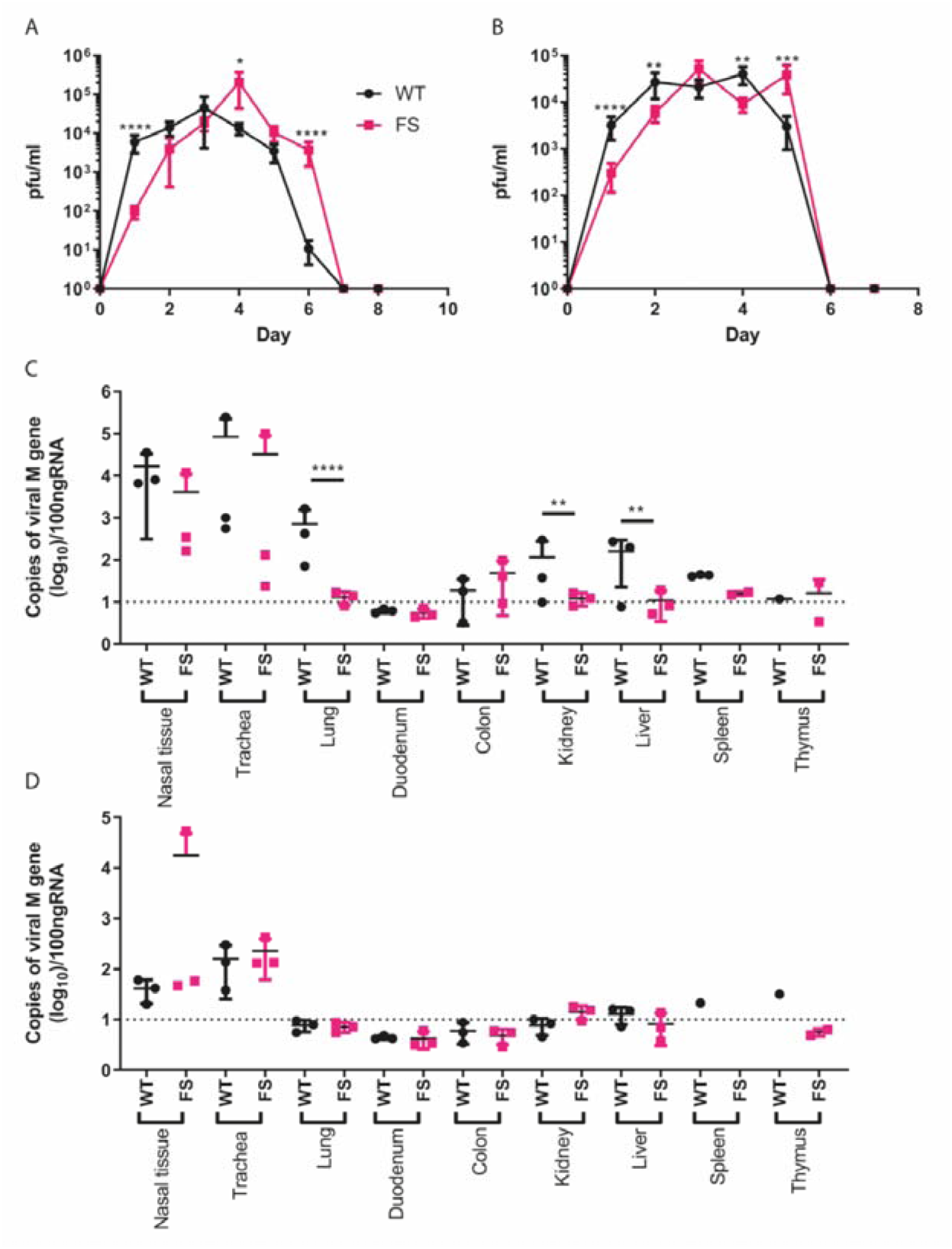
A lack of PA-X results in delayed shedding *in vivo* and restricted organ tropism. Birds were directly infected with either WT or FS H9N2 virus and naive contact birds were introduced one day post-inoculation. Swabs were taken from the buccal cavity throughout the study duration and virus titrated via plaque assay. The average +/− SD buccal shedding profile of at least 4 birds per group are shown for (A) directly infected birds and (B) contact birds. Statistics determined by Mann-Whitney U Test. (C, D) Detection of viral M gene within tissues of (C) directly infected and (D) contact birds. On day 2 post-infection 3 birds per group were culled, a panel of tissues taken and following RNA extraction qRT-PCR for viral M gene carried out. Dotted lines indicate limit of detection. CT values were compared to a M gene standard curve to determine copy number. Graphs show mean RNA copy number +/− SD. Statistics determined by unpaired T-test throughout. P values for statistics throughout: *, 0.05 ≥ P > 0.01; **, 0.01 ≥ P > 0.001; ***, 0.001 ≥ P > 0.0001; ****, P ≤ 0.0001.

All contact birds in both groups became infected and showed buccal shedding from 1 day post-exposure to the directly infected birds, indicating robust contact transmission of both viruses (Figure 4B). A similar buccal shedding pattern to the directly infected birds was seen in the contact birds; UDL-01 FS again showed delayed shedding kinetics, with significantly less virus shed on days 1-2 post-exposure and significantly more virus shed by day 5 post-exposure. Within both contact groups viral shedding was cleared by day 6 post-exposure. When AUC values were calculated for the contact bird populations, values were comparable; UDL-01 WT AUC was 94,171 whereas the UDL-01 FS AUC was 104,655. Overall, the shedding profiles of the infected birds suggested that expression of PA-X by the WT virus led to accelerated, but not increased, buccal shedding of virus compared to a virus which lacked PA-X expression.

On day 2 post-inoculation 3 birds from each group – directly infected, contact or mock – were euthanised and a panel of tissues were taken to assess viral tropism and cytokine profiles. After RNA extraction, qRT-PCR for the viral M gene was performed to assess viral replication within different tissues. In tissues isolated from directly infected birds, viral replication was primarily observed within the upper respiratory tract in both UDL-01 WT and UDL-01 FS infected birds (Figure 4C). There were no significant differences in M gene copy number within the nasal tissue and trachea. However, significantly more viral RNA was detected for UDL-01 WT in the lower respiratory tract in the lung. Within other visceral organs little viral RNA was detected, particularly in tissues collected from UDL-01 FS infected birds. UDL-01 WT RNA could be detected in both the liver and kidneys to significantly higher levels than UDL-01 FS. Therefore, UDL-01 WT virus showed increased viral dissemination compared to UDL-01 FS at day 2 post-inoculation. Tissues taken from the contact birds at day 2 of the experiment (i.e. day 1 post-exposure), again showed robust RNA levels in the upper respiratory tract tissues, the nasal tissue and trachea (Figure 4D). Viral loads were not significantly different between UDL-01 WT and FS infected birds in any tissues and very low levels of RNA were detected in non-respiratory tissues. Overall, these data show that removal of PA-X from UDL-01 led to reduced viral dissemination at day 2 post-infection for directly inoculated birds.

### Cytokine expression in the upper respiratory tract of directly infected birds correlated with viral titres

PA-X is known to alter host responses by downregulating protein synthesis (Jagger et al., 2012). Within different host species, modulation of PA-X expression has been shown to alter host expression of cytokines and chemokines; for example an H9N2 AIV unable to express PA-X has been shown to have decreased expression of *IL-6, IL-1β, CCL3, IFN-γ* and *TNF-α* within a mouse model compared to a virus with PA-X expression (Gao et al., 2015c). Therefore, it was assessed whether UDL-01 WT and FS led to differential cytokine and chemokine responses within the chicken host. The upper respiratory tract tissues were chosen due to robust and comparable viral replication in directly infected animals (Figure 4C). qRT-PCR for a range of chicken cytokines and markers of the interferon response were assessed. The host gene, *RPLPO-1*, was used for gene normalisation.

Nasal tissue from directly infected birds displayed little differences in immune response (Figure 5A). Some minor differences were seen with expression of *IFN-β, IFN-γ* and *IL-18* with UDL-01 WT infected birds generally expressing higher levels of cytokines. These differences only reached significance with the expression of *IL-1β* and the innate immune effector gene *Mx*, with UDL-01 WT infected birds expressing higher levels of these immune markers. Immune responses in the tracheas of directly infected birds showed a similar trend to those in the nasal tissue (Figure 5B). Few differences in cytokine expression were seen between UDL-01 WT and UDL-01 FS infected animals. UDL-01 WT infected animals again tended to have increased expression of *IFN-β, IFN-γ* and *IL-1β* although this only reached significantly different levels with *CXCLi2*. It was worth noting that although UDL-01 WT trended towards having higher levels of cytokines in the upper respiratory tract, these tissues also had higher levels of viral RNA, therefore it is difficult to draw conclusions about whether PA-X is having a direct role on cytokine expression; the trend for reduced cytokines in FS mutants may be a result of reduced viral RNA in tissues infected with these viruses.

**Figure 5.**
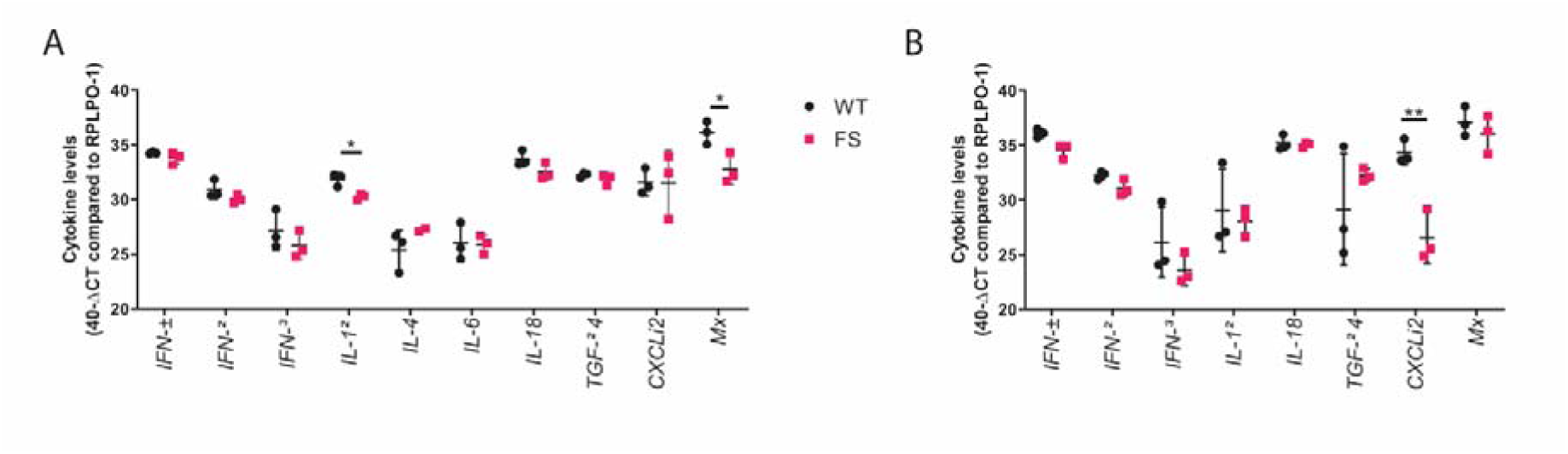
Cytokine induction in H9N2 infected birds. RNA was extracted from nasal tissue (A) or trachea (B) of WT and FS infected chickens. qRT-PCR was performed for a panel of cytokines and levels compared to the reference gene, RPLPO-1 were calculated. Cytokine expression for each bird is represented in a single point, with error bars displaying mean +/− SD of tissues from n=3 chickens. Statistics were determined by Unpaired T-tests. P values for statistics throughout: *, 0.05 ≥ P > 0.01; **, 0.01 ≥ P > 0.001

## Discussion

In this study we set out to investigate whether PA-X expression is a virulence factor for H9N2 viruses in their natural poultry host. As expected, ablating PA-X expression resulted in loss of host shutoff activity, showing that, as with other IAV strains, the shutoff activity of H9N2 viruses is partly due to PA-X expression. We found that PA-X expression resulted in slightly increased replication in mammalian MDCK cells but had no effect on titres in primary chicken cells or eggs, although virus with PA-X caused more embryonic lethality *in ovo* at higher input doses. We further found that *in vivo*, ablating PA-X expression led to a delayed shedding profile and lower visceral organ tropism. These results, the first study to investigate the impact of PA-X expression in a low pathogenicity avian influenza virus in its natural host suggest that PA-X expression aids faster virus replication and dissemination *in vivo*, although we saw little evidence for this acting through viral modulation of induced cytokine levels.

The finding here that expression of PA-X causes more rapid virus shedding after infection but that a virus with ablated PA-X expression potentially sheds for longer than WT virus is similar to what has been seen for H1N1-infected mice (Lee et al., 2017) and H9N2-infected mice (Gao et al., 2015c). The latter study showed that when mice were infected with a different lineage of H9N2 virus to the one used here (BJ94 versus G1) that had PA-X ablated, there was decreased virulence associated with reduced virus titres in mouse lungs. Interestingly, PA-X tells a contrasting story in high pathogenicity avian influenza viruses and the 1918 H1N1 pandemic virus. Loss of PA-X in 1918 H1N1 and HPAI H5N1 viruses caused increased virulence in mice (Jagger et al., 2012, Gao et al., 2015b, Gao et al., 2015a, Gao et al., 2015c) and in ducks and chickens (Hu et al., 2015). While Jagger and colleagues did not link the increased virulence in mice of 1918 H1N1 IAV lacking PA-X to effects on virus replication, Gao and colleagues showed that increased virulence in mice after loss of H5N1 PA-X was associated with increased titers of the PA-X null virus in the lungs, brain and blood of infected mice. Similarly, Hu and colleagues showed that increased virulence in chickens, ducks and mice on loss of H5N1 PA-X correlated with increased virus titers of the PA-X null virus.

The authors of the H9N2 infection study in mice (Gao et al. 2015c) proposed that differences in effects of PA-X on virulence could be due to the fact that high pathogenicity viruses induce high levels of cytokine responses but low pathogenicity viruses do not typically induce high levels of cytokines. Since a ΔPA-X mutant of a low pathogenicity virus is less effective at host cell shut off, it was more effective at eliciting an antiviral response resulting in reduced virus replication in their study.

Although no difference in pathogenicity in viruses expressing or lacking PA-X was seen in this study it should not be ruled out there may be a role. UDL-01 WT has shown to be a good model for a moderately pathogenic H9N2 virus in previous studies, causing clear clinical signs and even limited mortality (James et al., 2016, Peacock et al., 2017). However these studies used Rhode Island Red breed chickens, whereas here we used white Leghorn birds which appear to be more resilient to avian influenza virus infection (Sironi et al., 2008, Matsuu et al., 2016). Therefore, it is possible that the virus lacking PA-X is less pathogenic than UDL-01 WT, but this was not observed due to the lack of clinical signs and mortality in the UDL-01 WT infected groups in this system. It is worth noting we have previously correlated visceral tropism with pathogenicity and clinical signs in UDL-01 (James et al., 2016, Peacock et al., 2017); in this study UDL-01 WT did show greater visceral tropism than UDL-01 FS, perhaps suggesting an attenuated pathogenicity when PA-X expression is ablated.

Overall, this work suggests PA-X may play a role in H9N2 viruses in birds by allowing more rapid replication and dissemination throughout the host, potentially leading to higher pathogenicity. This work will be useful in future surveillance efforts allowing the assessment of newly sequenced viruses as it suggests viruses expressing a full-length PA-X are likely to have a wider tropism and higher pathogenicity than those that do not. Furthermore, this work suggests that there may be slightly different roles for PA-X in mammalian and avian hosts, potentially helping explain the mechanism by which PA-X works.

## Materials and methods

### Ethics Statement

All animal experiments were carried out in strict accordance with the European and United Kingdom Home Office Regulations and the Animal (Scientific Procedures) Act 1986 Amendment regulation 2012, under the authority of a United Kingdom Home Office Licence (Project Licence Numbers: P68D44CF4 X and PPL3002952).

### Cells

Madin-Darby canine kidney (MDCK) cells, human embryonic kidney (HEK) 293T cells and chicken DF-1 cells were maintained in Dulbecco’s Modified Eagle Medium (DMEM; Sigma) supplemented with 10% FBS and 100 U/ml Penicillin-Streptomycin (complete DMEM). All cells were maintained at 37°C, 5% CO_2_.

Primary chicken kidney (CK) cells were generated as described elsewhere (Hennion and Hill, 2015). Briefly, kidneys from three-week-old SPF Rhode Island Red birds were mechanically shredded, washed, trypsinised, and then filtered. Cells were resuspended in CK growth media (EMEM + 0.6% BSA, 10% v/v tryptose phosphate broth, 300 U/ml penicillin/streptomycin), plated and maintained at 37°C, 5% CO2.

### Viruses

All work in this study was performed with the reverse genetics derived H9N2 virus, chicken/Pakistan/UDL-01/2008 (UDL-01)(Long et al., 2016). UDL-01 virus segments were expresd in the bi-directional reverse genetics PHW2000 plasmids (Hoffmann et al., 2000). Mutant PA segments were generated by site directed mutagenesis.

Viruses were rescued as described elsewhere (Hoffmann et al., 2000). Briefly, 250ng of each segment plasmid were co-transfected into 293Ts plated in 6 well plates using lipofectamine 2000. 16h post-transfection, media was changed to reverse genetics media (DMEM + 2mM glutamine, 100U/ml penicillin, 100U/ml streptomycin, 0.14% BSA, 5µg/ml TPCK-treated trypsin). Following a 48h incubation at 37°C, 5% CO_2_, supernatants were collected and inoculated into embryonated hens’ eggs for subsequent harvest of virus infected allantoic fluid.

### *In vitro* translation and autoradiography

*in vitro* translations to visualise radiolabelled proteins, the TnT® Coupled Reticulocyte Lysate system (Promega; #L4610) was utilised as per the manufacturers’ instructions using 200ng of plasmid DNA. Reactions were incubated at 37°C for 90 min then denatured in 50µl protein loading buffer and analysed via SDS-PAGE and autoradiography. X-ray films were developed using a Konica SRX-101A X-ograph film processor as per the manufacturers’ instructions.

### Host cell shutoff assays

β-gal shutoff reporter assays were performed as described elsewhere (Jagger et al., 2012). Briefly, 293T or DF-1 cells were co-transfected with expression plasmids for the influenza segment 3 (PA) and β-galactosidase (β-gal) reporter. 48 hours later, cells were lysed with 100 µl of 1x Reporter lysis buffer (Promega). β-gal expression was assayed using the β-galactosidase enzyme assay system (Promega). A Promega GloMax Multi Detection unit was used to measure absorbance at 420 nm.

For the shutoff activity assays using live virus, MDCKs were infected with virus at a high MOI of 5. At 7.5 hours post-infection, cells were washed and the medium changed to 1ml of complete DMEM containing 10 µg/ml of Puromycin dihydrochloride from *Streptomyces alboniger* for 30 min. Cells were washed and lysed in protein loading buffer and run on an SDS-PAGE and were then western blotted, probing for puromycin. Puromycylated protein synthesis was quantified in the region of the gel between 45 kDa and 80 kDa. Protein quantification following western blot was measured by densitometry using ImageJ analysis software.

### Mini-replicon assays

293T or DF-1 cells were transfected with plasmids encoding PB2, PB1, PA and NP, along with a firefly luciferase vRNA-like reporter under a cell-type specific polI promoter (human for 293T, chicken for DF1). The following concentration of plasmid were used for 293Ts, PB2-80 ng, PB1-80ng, PA-20 ng, NP-160 ng, pPol I Luc-800 ng, these amounts were doubled for DF1s. After 48 hours, media was removed, and cells were lysed in 100 µl of 1x Passive Lysis Buffer (Promega). A Promega GloMax Multi Detection unit was used to measure luciferase activity following the manufacturer’s instructions.

### Virus replication assays

MDCK and CK cells were infected at a low MOI of 0.01 for 1 hour in serum-free DMEM, after which media was replaced with DMEM, 2 μg/ml tosyl phenylalanyl chloromethyl ketone (TPCK)-treated trypsin (MDCK cells) or Eagle’s minimum essential medium (EMEM), 7% bovine serum albumin [BSA], and 10% tryptose phosphate broth (CKs). Time points were harvested in triplicate at 4-, 8-, 12-, 24-, 48- and 72-hours post-infection. Virus titres were determined by plaque assay on MDCK cells.

10-day old embryonated hens’ eggs (VALO breed) were inoculated with 100 pfu of diluted virus into the allantoic cavity. Eggs were incubated for 4-72 hours and culled via the schedule one method of refrigeration at 4°C for a minimum of 6 hours. 5 eggs were used per virus per time point. Harvested allantoic fluid from each egg was collected and virus titres were assessed by plaque assay on MDCK cells.

For the egg mortality rates experiment a 10-fold serial dilution of each virus (10000 pfu to 10 pfu) was made as used to infect 5 embryonated eggs per virus per dilution. Embryos were candled twice daily throughout the study period to check for embryo viability (up to 84 hours post-infection). If eggs reached a predetermined end point at candling, they were deemed to be dead and the eggs chilled to ensure death before disposal. Markers of the end point included, a lack of movement of the embryo, disruption of blood vessels within the egg and/or signs of haemorrhage. If the embryos survived until the experimental end (84 hours post-infection) they were culled via a schedule one method and samples of allantoic fluid were collected to determine presence of virus via immunostaining for viral NP protein. Any eggs without positive detection of viral NP were removed from the study.

### Virus infection, transmission and clinical outcome *in vivo*

In vivo studies were performed with three-week-old White Leghorn birds (VALO breed). Prior to the start of the experiments, birds were swabbed and bled to confirm they were naïve to the virus. All infection experiments were performed in self-contained BioFlex B50 Rigid Body Poultry isolators (Bell Isolation Systems) at negative pressure. 10 birds per group were directly inoculated with 10^4^ pfu of virus via the intranasal route. Mock infected birds were inoculated with sterile PBS as an alternative. One day post-inoculation 8 naïve contact birds were introduced into each isolator to determine viral transmissibility.

Throughout the experiment, birds were swabbed in the buccal and cloacal cavities (on day 1-8, 10 and 14 post-infection). Swabs were collected into 1ml of virus transport media (WHO standard). Swabs were soaked in media and vortexed for 10 seconds before centrifugation. Viral titres in swabs were determined via plaque assay.

At day 2 post-infection, 3 birds per group (directly infected, contact and mock infected) were euthanised via overdose of pentobarbital (at least 1 ml). A panel of tissues were collected and stored in RNA later at −80°C until further processing. Birds were observed twice daily by members of animal services and whilst procedures were carried out for the presence of clinical signs of infection. On day 14 post-infection, all remaining birds were culled via overdose of pentobarbital or cervical dislocation.

### RNA extraction and RT-PCR from chicken tissues

30mg of tissue collected in RNA later was mixed with 750 µl of Trizol. Tissues were homogenised using the Retsch MM 300 Bead Mill system (20 Hz, 4 minutes). 200 µl of chloroform was added per tube, shaken vigorously and incubated for 5 min at room temperature. Samples were centrifuged (9,200 x g, 30 minutes, 4°C) and the top aqueous phase containing total RNA was added to a new microcentrifuge tube, subsequent RNA extraction was then carried out using the QIAGEN RNeasy mini kit following manufacturers’ instructions.

100 ng of RNA extracted from tissue samples was used for qRT-PCR. All qRT-PCR was completed using the Superscript III platinum One-step qRT-PCR kit (Life Technologies) following manufacturer’s instructions for reaction set up. Cycling conditions were as follows: i) 5 minutes hold step at 50°C, ii) a 2 minute hold step at 95°C, and 40 cycles of iii) 3 seconds at 95 °C and iv) 30 second annealing and extension at 60 °C. Cycle threshold (CT) values were obtained using 7500 software v2.3. Mean CT values were calculated from triplicate data. Negative controls were included within each plate to determine any unspecific amplification or contamination. Within viral M segment qRT-PCR an M segment RNA standard curve was completed alongside the samples to quantify the amount of M gene RNA within the sample from the CT value. T7 RNA polymerase-derived transcripts from UDL-01 segment 7 were used for the preparation of the standard curve.

Within the cytokine qRT-PCRs, three housekeeping genes were included per sample (RPLPO-1, RPL13 and 28S rRNA) that had been previously determined to be stable in a broad range of tissues. Briefly, the geNorm algorithm (Vandesompele et al., 2002) was adopted to calculate the stability for each reference gene and the optimal reference gene number from raw Cq values of candidate reference genes using qbase+ real-time qPCR software version 3.0 (Biogazelle).

## Acknowledgments

We would like to thank the animal housing staff for looking after the wellbeing of chickens used in this study and for monitoring their health throughout the experiments.

